# Docking-based long timescale simulation of cell-size protein systems at atomic resolution

**DOI:** 10.1101/2022.06.07.495183

**Authors:** Ilya A. Vakser, Sergei Grudinin, Nathan W. Jenkins, Petras J. Kundrotas, Eric J. Deeds

## Abstract

Computational methodologies are increasingly addressing modeling of the whole cell at the molecular level. Proteins and their interactions are the key component of cellular processes. Techniques for modeling protein interactions, so far, have included protein docking and molecular simulation. The latter approaches account for the dynamics of the interactions, but are relatively slow, if carried out at all-atom resolution, or are significantly coarse-grained. Protein docking algorithms are far more efficient in sampling spatial coordinates. However, they do not account for the kinetics of the association (i.e., they do not involve the time coordinate). Our proof-of-concept study bridges the two modeling approaches, developing an approach that can reach unprecedented simulation timescales at all-atom resolution. The global intermolecular energy landscape of a large system of proteins was mapped by the pairwise Fast Fourier Transform docking and sampled in space and time by Monte Carlo simulations. The simulation protocol was parametrized on existing data and validated on a number of observations from experiments and molecular dynamics simulations. The simulation protocol performed consistently across very different systems of proteins at different protein concentrations. It recapitulated data on the previously observed protein diffusion rates and aggregation. The speed of calculation allows reaching second-long trajectories of protein systems that approach the size of the cells, at atomic resolution.

## INTRODUCTION

Rapid progress in experimental and computational techniques is redrawing the map of molecular and cellular biology, eliminating old boundaries between research fields, and creating new opportunities for breakthroughs. In structural biology, AlphaFold has achieved unprecedented near-experimental accuracy in predicting the structure of individual proteins^1^ and, at the same time, is successfully using a similar approach in a different research field - protein docking - to predict the structure of protein complexes.^2^ Techniques for modeling protein interactions, so far, have consisted of two major categories: (1) protein docking,^3^ such as the Fast Fourier Transform (FFT) algorithm (which in short computing times performs full systematic search through translational and rotational degrees of freedom),^4^ and (2) molecular simulations, such as Molecular Dynamics (MD) or Brownian Dynamics (BD). Borrowing from the 4D space-time continuum terminology, protein docking has been restricted to sampling of the intermolecular energy landscape at atomic resolution in the 3D space component only, whereas atomic resolution molecular simulation protocols sample the entire 4D landscape albeit, due to the high computational cost, for short timescales only. Simulation approaches have been applied before, across the fields, to the protein docking problem, broadly for the refinement of the docking global search predictions,^5^ with more advanced approaches addressing the global docking search itself.^6–8^ Our study puts forward the reverse across-the-fields application of the docking techniques to the dynamics of the protein interactions.

The great accomplishments in structure prediction based on the deep learning do not solve the protein docking problem. This problem, traditionally thought of as a 3D one, simply requires adding the missing time coordinate from the docking space-time continuum. Re-focusing docking from the problem of finding the unique global minimum solution, to sampling the enormous multitude of transient interactions^9^ dominating the crowded cellular environment, allows propagating protein interactions in time. Such propagation can take full advantage of the vast amount of powerful and efficient methodologies accumulated in the protein docking field.^3^ Thus, it opens extraordinary new opportunities in structural modeling of the biomolecular mechanisms, allowing modeling of larger systems, at longer timescales, all based on the inherent to docking atomic resolution.

In the context of the spectacular advances in experimental and computational structural biology, structure-based modeling of protein interactions in the living cell is becoming more central than ever before.^10–12^ Traditional simulation protocols (such as MD and BD) are either relatively slow, if carried out at the all-atom representation,^13^ or significantly coarse-grained, with one particle representing a protein.^14^ Thus, there are only a few examples of structure-based simulations at the scale of the whole cell.^11, 13, 15^ Cell modeling is important for a variety of reasons, including integration of data into a unified representation of knowledge about an organism, prediction of multi-network phenotypes, filling the gaps in our knowledge of cellular processes, and development of our ability to modulate them.^10, 16–18^ Early approaches to cell modeling represented proteins by hard spheres.^14, 19^ BD simulations of a part of the *E. coli* cytoplasm were run for 20 μs in rigid body all-atom representation,^20^ coarse grained in a subsequent study.^21^ All-atom MD simulations of bacterial cytoplasm were run for 100 ns.^22^ Since then, the all-atom MD simulations of cellular environment reached the μs timescales.^13, 23–25^ Modeling also has been used to study the confinement effect and hydrodynamic properties of the crowded environment,^26^ the physical limits of cells,^27^ and packing of the cellular environment.^28, 29^ The FFT approach was used to study protein folding and binding in the crowded environment^30, 31^ and in the free energy calculations.^32^

It has been commonly accepted that mesoscopic particles, such as proteins, in simple solvents can be described with Brownian diffusion. However, this description fails dramatically with molecules in complex biological media, such as the cellular environment.^33, 34^ While theoretical models can, in principle, explain some of these effects, their applicability requires *a priori* knowledge of the molecular organization of crowding particles in time and space.^35^ Thus, simulation techniques, such as MD or BD, are currently the only computational way to access dynamical characteristics of cellular environments. MD simulations are usually restricted to very short time scales. BD simulations allow access to much longer times, but require careful mesoscopic parameterization, e.g., with diffusion constants. An alternative to these simulation methods is Monte Carlo (MC) protocols, which allow computing kinetic parameters, such as diffusion coefficients. It requires only computation of the system’s potential energy at each time step. MC estimate of the self-diffusion coefficient in the continuous move case is in good agreement with the BD simulations.^36^

Rigorous experimental tests of the predictions from cell simulations have remained elusive. They have focused almost exclusively on validating predictions of the diffusion coefficient of a protein in a crowded cellular environment by measurements of fluorescent proteins diffusion in cells.^10, 22^ These results showed that effects like transient interactions and excluded volume significantly decrease the rate of diffusion of proteins in cells.^10^

Our proof-of-concept study linked FFT-accelerated systematic docking with the MC simulations, allowing propagation of large protein systems in time with great computational efficiency. The approach was validated on experimental and computational observations from prior studies and is capable of reaching second-long simulations of the cellular environment at all-atom resolution.

## METHODS

### Modeling paradigm

Our approach was to dramatically speed-up the sampling of the intermolecular energy landscape by skipping the low-probability (high-energy) states, focusing only on the set of high-probability (low-energy) states corresponding to the energy minima. The “minima hopping” paradigm has been widely used since the early days of molecular modeling for the sampling of the energy landscapes of biomolecules - such as conformational analysis of biopolymers,^37^ rotamer libraries,^38^ and refinement of protein-protein interfaces,^39^ providing extraordinary savings of computing time by avoiding travel in low-probability areas of the landscape. Markov State Models (MSM), have been used to study protein folding, dynamics,^40^ and association^41^ by representing the energy landscape by a set of the energy minima and the probabilities of transition between them. In this study we use a similar idea, namely a Markov State Monte Carlo approach to sampling transitions between low energy states, to perform very long trajectory simulations of large systems of proteins at atomic resolution.

### Molecular systems

Simulations were performed on three different sets of proteins. To determine the volume fraction of the system, for each protein, the volume was calculated by the 3V server (http://3vee.molmovdb.org).^42^

*Set 1*. Five arbitrarily selected globular proteins of average size to represent a “typical” crowded cellular environment (hereafter called “5 mix” set; Fig. 1A and Table 1).

**Figure 1.**
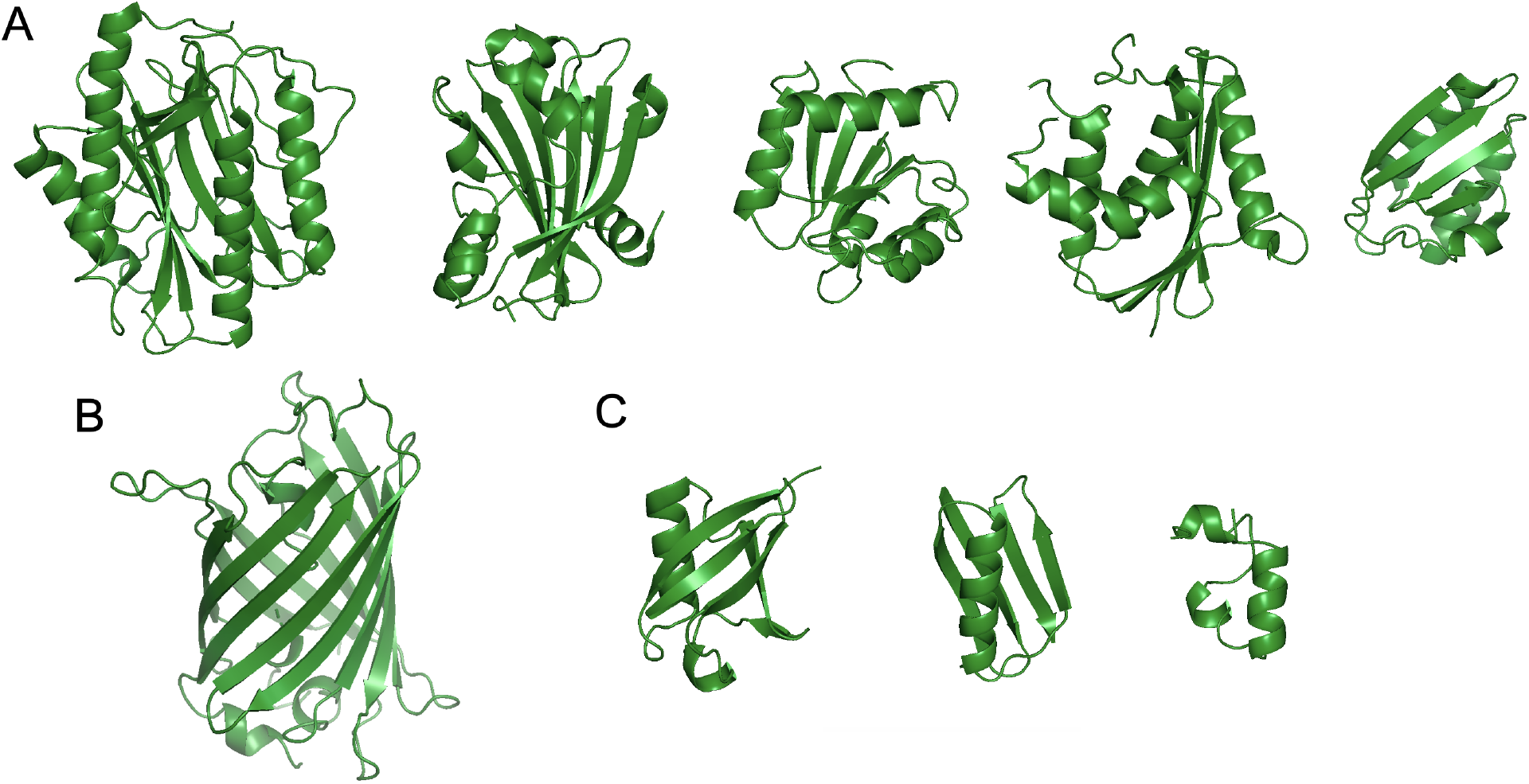
Molecular systems used in the study. (A) Five arbitrarily selected globular proteins of average size to represent a typical crowded cellular environment (PDB codes 1mat, 1g81, 3chy, 1jxb and 1cm2). (B) Green Fluorescent Protein (1ema). (C) Three small proteins: ubiquitin (1ubq), G-protein B subunit (1pga) and villin (1vii). Molecular images were obtained using PyMOL.^50^

**Table 1.** Characteristics of the proteins.

| Size | 5 mix set |  |  |  |  | GFP | 3 mix set |  |  |
| --- | --- | --- | --- | --- | --- | --- | --- | --- | --- |
|  | 1mat<sup>1</sup> | 1g81 | 3chy | 1jxb | 1cm2 | 1ema | 1ubq | 1pga | 1vii |
| No. of residues | 263 | 163 | 128 | 152 | 85 | 210 | 76 | 56 | 35 |
| Volume, Å<sup>3</sup> | 40,693 | 27,823 | 21,151 | 25,753 | 13,720 | 36,767 | 12,965 | 9,019 | 6,583 |

<sup>1</sup>PDB codes.

*Set 2*. Set 1 plus Green Fluorescent Protein (“GFP + 5 mix” set; Fig. 1B and Table 1) for comparison with the experimental data on GFP diffusion.

*Set 3*. Three small proteins (“3 mix” set; Fig. 1C and Table 1) from Feig and co-workers^15^ representing the non-membrane part of that study: Ubiquitin, G-protein B subunit, and Villin.

### Generation of the initial state

For the starting point of the simulation, the proteins were placed on a cubical grid of a pre-set size, with the step of the grid calculated according to the desired protein volume fraction. In this study, we used 500 x 500 x 500 Å^3^ grid (the linear dimension about half of that of the smallest cell - *Mycoplasma genitalium*) with periodic boundary conditions. Each protein had an equal share of copies (e.g., in the “5 mix” set of the five proteins mixture, each protein had 1/5 share of copies). The total number of protein copies and the step of the grid were calculated according to the pre-set protein volume fraction *V*. In this study, we used a range of volume fraction values, from *V* = 0.10 to close to physiological *V* = 0.30. Table 2 shows the total number of molecular copies corresponding to each volume fraction.

**Table 2.** Characteristics of the molecular systems.

| Volume fraction | Number of molecules in simulation system |  |  |
| --- | --- | --- | --- |
|  | 5 mix set | GFP + 5 mix<sup>1</sup> | 3 mix set<sup>2</sup> |
| 0.10 | 460 |  | 1,314 |
| 0.15 | 725 |  |  |
| 0.20 | 970 |  |  |
| 0.25 | 1,210 |  |  |
| 0.30 | 1,450 | 1,356 | 3,939 |

<sup>1</sup>GFP + 5 mix set was run only at physiological volume fraction 0.30 at which the experimental data was obtained.
<sup>2</sup>3 mix set was run only at volume fractions 0.1 and 0.3 for which the molecular dynamics data was available.

The proteins were placed in a random order. They were randomly rotated and translated within half of the grid step interval. No collision check was applied at this stage since the collisions were eliminated at the start of the simulation. Supplementary Information Figure S1 shows a fragment of the initial state of the system before the start of the simulation.

### Simulation protocol

An MC procedure was developed to simulate the cellular environment with proteins in rigid-body approximation, using an all-atom representation. The procedure by design is based on proteins transitioning between different protein-protein associations. Thus, our approach applies to crowded protein environments only, where proteins encounter each other in close proximity, and monomeric states (the absence of all protein-protein interactions, including transient) are uncommon. The energy landscape of the system was approximated by our GRAMM FFT docking^4, 43^ scores/energies, in which the docking poses (including the multiplicity of transient encounters) corresponded to negative energy values, and the monomeric states (i.e., the barriers between the minima) have energy zero.

The position of each protein was described with the 3 x 3 rotation matrix and the translation vector relative to the origin of the coordinate system. Protein-protein docking poses were systematically pre-computed for all rotations and translations of each protein, relative to all other proteins in the system by GRAMM docking, unscored and unrefined, at intermediate resolution, previously optimized for the docking of unbound proteins^44^ (grid step 3.5 Å, repulsion 9.0, and rotation interval 10°). For proteins A and B, both docking combinations A - B (A is the ligand, and B is the receptor) and B - A (B - ligand, A - receptor) were precalculated. Thus, e.g., for the “5 set” the number of precalculated docking outputs was 25 (5 x 5). If A was the moving molecule (ligand), its new putative energy was taken from the A - B docking (and vice versa).

The docking results were stored on six-dimensional grids (three translations and three rotations) and accessed during the MC runs. The MC move set included a move of one randomly selected protein (“ligand”) to another randomly selected protein (“receptor”) within a certain neighborhood (described below) from the ligand’s current position. Once the ligand and the intended receptor are selected, the move is chosen randomly among the precalculated top 30,000 docking matches for that pair of proteins.

The simulation step is completed when all proteins have attempted to move. Once the ligand moves, the energy (GRAMM docking score) of the new match is added to the ligand’s energy and the energy of the old match it detaches from is subtracted. Correspondingly, its new receptor’s energy adds that new match’s energy, and the old receptor’s (the one the ligand is detaching from) energy subtracts the energy of the detaching docking match.

The move is accepted or rejected based on the Metropolis acceptance criterion (detailed balance condition). Ligands (L) are allowed to move to the neighboring receptors (R) only, defined as those within the distance between R and L geometric centers less than the sum of the R and L radii, plus 50 Å, to accommodate binding to the first layer of receptors in the crowded environment. Collision check is performed for each attempted move according to C^α^ - C^α^ minimal distance of 8 Å. The moves resulting in collision are rejected. Figure 2 illustrates the general principle of the move set. Periodic boundary conditions were introduced. Temperature was a parameter to be adjusted for an adequate acceptance rate.

**Figure 2.**
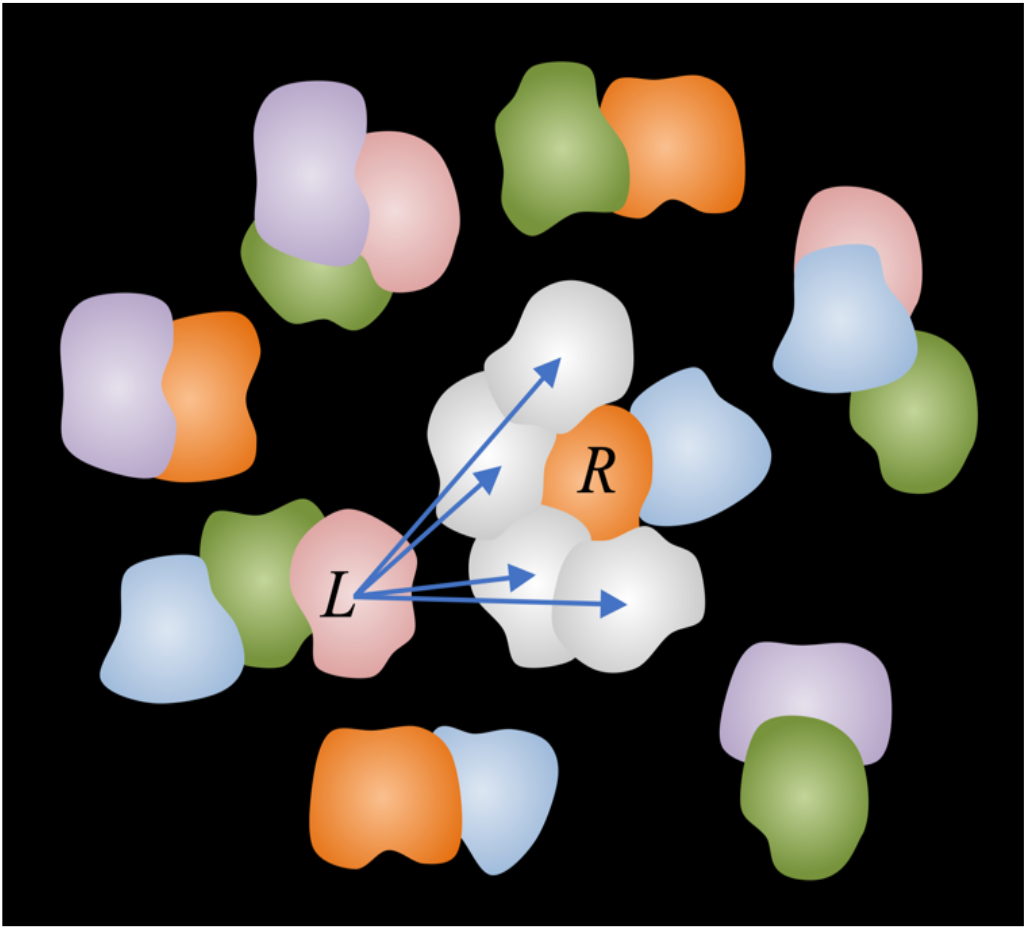
The simulation move set. Docking results are precalculated and stored on six-dimensional grids, accessed during the MC runs. The move set includes a move of one protein (*L* - ligand) at a time to a putative docking match with another protein (*R* - receptor) in the vicinity of the ligand. The energies of the states are set according to the docking scores. The move is accepted or rejected based on the Metropolis criterion (detailed balance condition).

The detailed balance condition for the system was implemented. The probability *P_ij_* of move from step *i* to step *j* had to be the same as *P_ji_* from *j* to *i*. Accordingly, the Metropolis criterion was normalized^45^ as

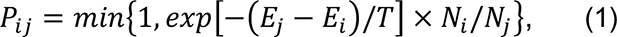

where *N_m_* is the numbers of possible moves (receptors to move to; Fig. S2) from state *m* with probability to be selected 1/*N_m_*; *E_m_* is the energy of state *m*; and *T* is the temperature (a scaling factor).

As noted above, in our system, the monomeric states have energy zero, and all minima have negative energy values. Our model assumes no additional barriers between states *i* and *j*. We also assume the same curvature of the potential wells of each state. Thus, in the Kramers (or Arrhenius) rate equation, which for our system can be written as

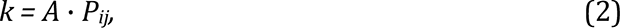

where *k* is the rate constant and *P_ij_* is the energy and temperature-based probability of move from step *i* to step *j* (Eq. 1), the pre-factor *A* is the same for all transitions. Thus, our scheme differs from the Kinetic Monte Carlo, because the transition rates are computed on-the-fly at each step and are proportional (with the constant *A*) to the acceptance probability of a new state.

Observed parameters of the simulation (per simulation step) were: potential energy *E* - the average energy of a molecule (the sum of all molecules’ energies - GRAMM docking scores - divided by the number of molecules); the shift (the average length of a molecule’s move per simulation step); the MSD (the average mean square deviation of a molecule’s geometric center after unwrapping coordinates from the periodic boundary conditions); acceptance rate (percentage of accepted moves), and the aggregation number *N_c_* (the average number of proteins in an aggregate/oligomer formed by docked proteins). To allow off-the-grid relaxation of the system, the reference position for MSD calculation was set at step 100. Diffusion rates *D_t_* were calculated from the slope of MSD according to the Einstein relationship *D_t_* = MSD(*t*)/6*t*, where *t* is the lag time.

## RESULTS AND DISCUSSION

### Temperature

The results of the simulation on the “5 mix” set at the physiological volume fraction (Fig. 3) and lower volume fractions (Fig. S3) showed that at low temperatures, the system is frozen (little to no movement of the proteins). At high temperatures, the system is overheated (moves accepted regardless of the energy). The melting curves (Fig. 3 and Fig. S3) had a clear inflection point at *T* = 100, consistently at all volume fractions, at which the system melts (breaks from the freeze) but is not overheated yet, and thus is likely most representative of the physiological conditions. The value of *T* corresponding to the melting phase transition reflects the docking energy landscape (mapped in GRAMM energy units), as follows from Eq. 1, namely the energy gap between a few deep minima (frozen system states) and multiple high-energy/transient states (melted system).

**Figure 3.**
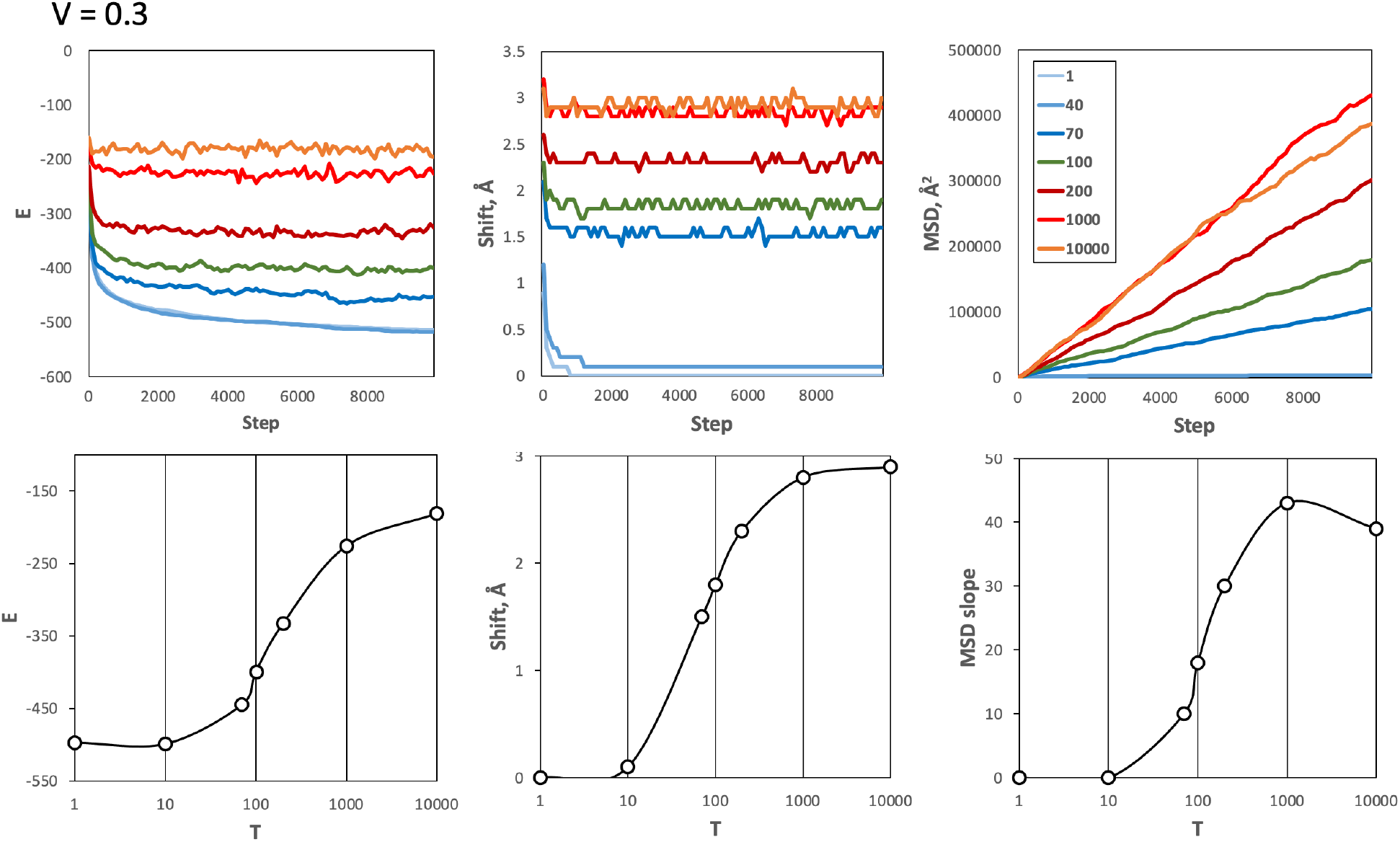
Simulations of the “5 mix” set at physiological volume fraction and a range of temperatures. The volume fraction *V* was set to close to physiological 0.3 value. The top panels show the energy *E*, shift, and MSD vs. simulation steps. MSD was calculated as the average for 1mat proteins. The temperatures *T* = 1 - 10,000 are shown by different colors. The data on the plots was smoothed by a 100-steps averaging sliding window. At low temperatures, the system is frozen (little or no movement of the proteins). At high temperatures, the system is overheated (moves accepted regardless of the energy). The melting curves (the bottom panels in log scale) have a clear inflection point at *T* = 100 indicating the optimal temperature at which the system melts (breaks from the freeze) but is not overheated yet.

Simulation on the “3 mix” set, which is a very different system from the “5 mix” set (the “3 mix” proteins are much smaller than the ones in the “5 mix”) yielded virtually identical melting behavior, at all volume fractions, with the same optimal temperature *T* = 100 (Fig. S4). This confirms the robustness of our approach and adds evidence to the validity of our approximation. Accordingly, for the rest of this study, we used *T* = 100 as the temperature of the systems.

### Calibration

We calibrated the time units of the simulation protocol on the available data from MD simulation of Villin at the physiological volume fraction in the non-membrane system.^15^ Here, the diffusion coefficient *D_t_* value was determined to be 3.5 Å^2^/ns, which according to the authors is three times greater than in experiment. Our simulation of the Villin within the “3 mix” protein set at the physiological volume fraction (Fig. 4) allowed us to calibrate our system’s time variable *t*, by matching the *D_t_* values calculated as *D_t_* = MSD/6*t* (see Methods) with the MD results, corrected by the above-mentioned factor of 3. Accordingly, one step of our simulation protocol was determined to be 20 ns.

**Figure 4.**
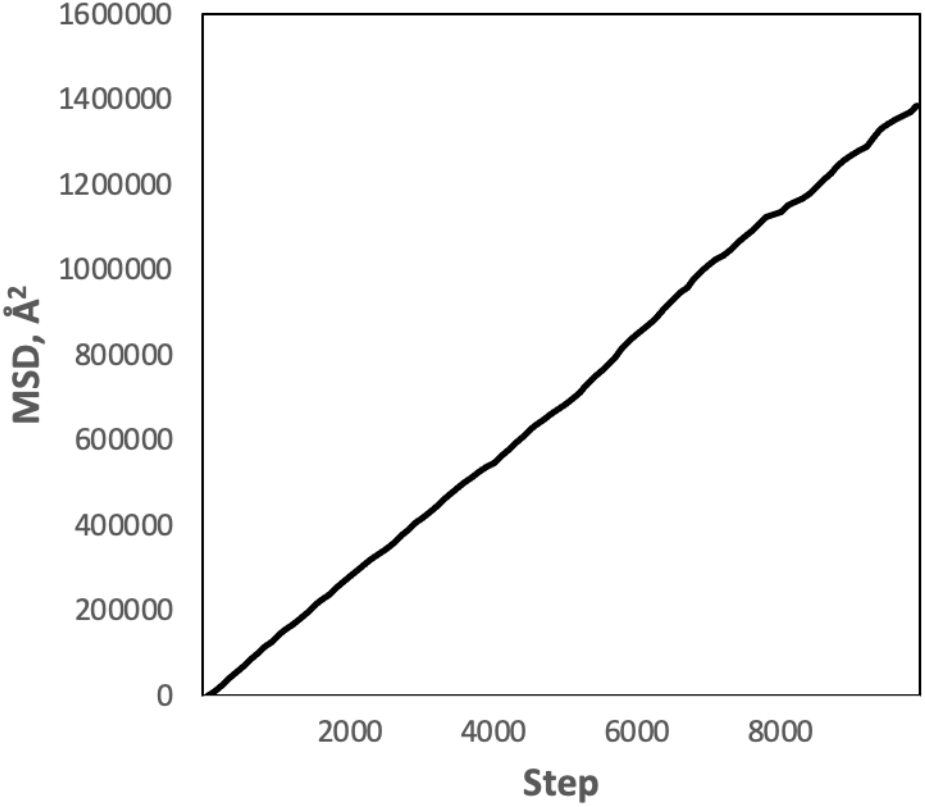
Simulation of Villin within the “3 mix” protein set. The simulation was run at *T* = 100 and *V* = 0.3 (see text). MSD was calculated as the average for the Villin proteins. The details of the observable parameters are the same as in Figure 3. The system’s time variable *t* was calibrated by matching the *D_t_* value, calculated from the slope of the MSD, as *D_t_* = MSD/6*t*, with the previously determined *D_t_* values (see text). One step of our simulation protocol was thus determined to be 20 ns.

### Validation and Quantitative Characterization of Protein Systems

The simulation protocol was validated on a number of observable parameters, testing for consistency of the results and correspondence to experimental and modeling studies. Our “minima hopping” paradigm, which by design allows no intermediate states between the minima (the minima correspond to the protein bound to another protein), assumes close proximity of the minima to each other (i.e., a crowded environment). Thus, our approximation would not hold for dilute systems. However, it allows for an observation of quantitative characteristics at a range of volume fractions. In our study, this range was set from 0.1 to close to physiological 0.3.

#### Melting temperature

As described above, the melting temperature for very different protein systems - the “5 mix” set of average size proteins and the “3 set” of much smaller proteins - at the full range of volume fractions, from 0.1 to 0.3, is the same. This supports the validity of our approximation and its consistency across different concentrations and size scales of proteins.

#### Diffusion rate in different systems

Experimental data on the diffusion of GFP in the cytoplasm of *Escherichia coli*^46^ puts the GFP diffusion coefficient *D_t_* in the 0.2 - 0.9 Å^2^/ns range. We ran simulation of the GFP with the “5 mix” protein set at a physiological volume fraction. The results (Fig. 5) showed the GFP diffusion rate was 0.3 Å^2^/ns, in excellent agreement with the experiment. It provides another confirmation of the approach validity and consistency across very disparate systems of proteins.

**Figure 5.**
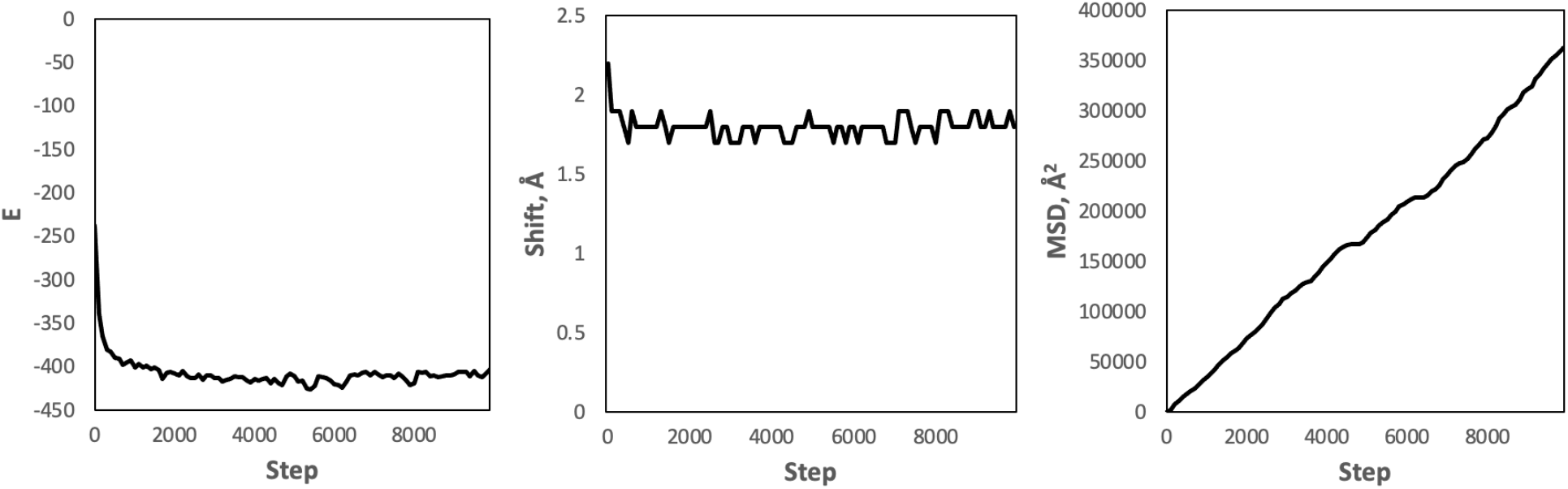
Simulation of the GFP with the “5 mix” protein set. The simulation was run at *T* = 100 and *V* = 0.3 (see text). MSD was calculated as the average for the GFP proteins. The details of the observable parameters are the same as in Figure 3.

#### Diffusion rate dependence on concentration

Simulation in the “5 mix” set at different volume fractions showed a pronounced slowdown of the diffusion with the increase of the protein concentration (Fig. 6) in accordance with long established evidence.^13, 15^ The quantitative scope of this slowdown (the ratio of the diffusion rates), for our range of volume fractions, is available from the MD simulation for Villin as 5.4, from *V* = 0.1 to 0.3.^15^ In our simulation of the “3 mix” set, that slowdown for Villin was 3.4, in good agreement with the MD data.

**Figure 6.**
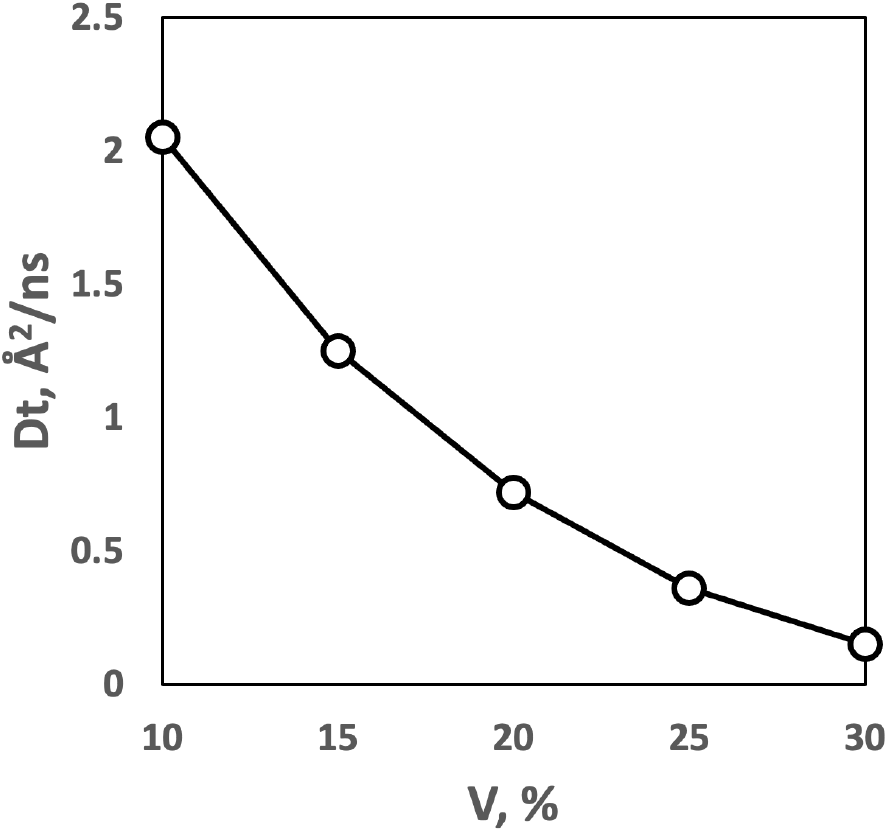
The slowdown of protein diffusion with the increase of protein volume fraction. D_t_ was calculated for 1mat proteins in the “5 mix” set.

#### Diffusion rate dependence on size

It is well established by experiment and simulation that larger proteins diffuse at a slower rate.^15, 46^ Due to the complexity and heterogeneity of the systems, the quantitative estimates of the size *vs*. diffusion correlation vary significantly. Our simulation of sets of small proteins *vs*. those of much larger proteins (see above) showed that the smaller ones diffuse significantly faster. Diffusion of proteins in the same “5 mix” set simulation showed clear size vs. diffusion rate correlation, at all volume fractions. The rate of the slowdown grows significantly with the increase of the volume fraction (Fig. 7). A similar trend was observed in the simulation of the “3 mix” set (Fig. S5).

**Figure 7.**
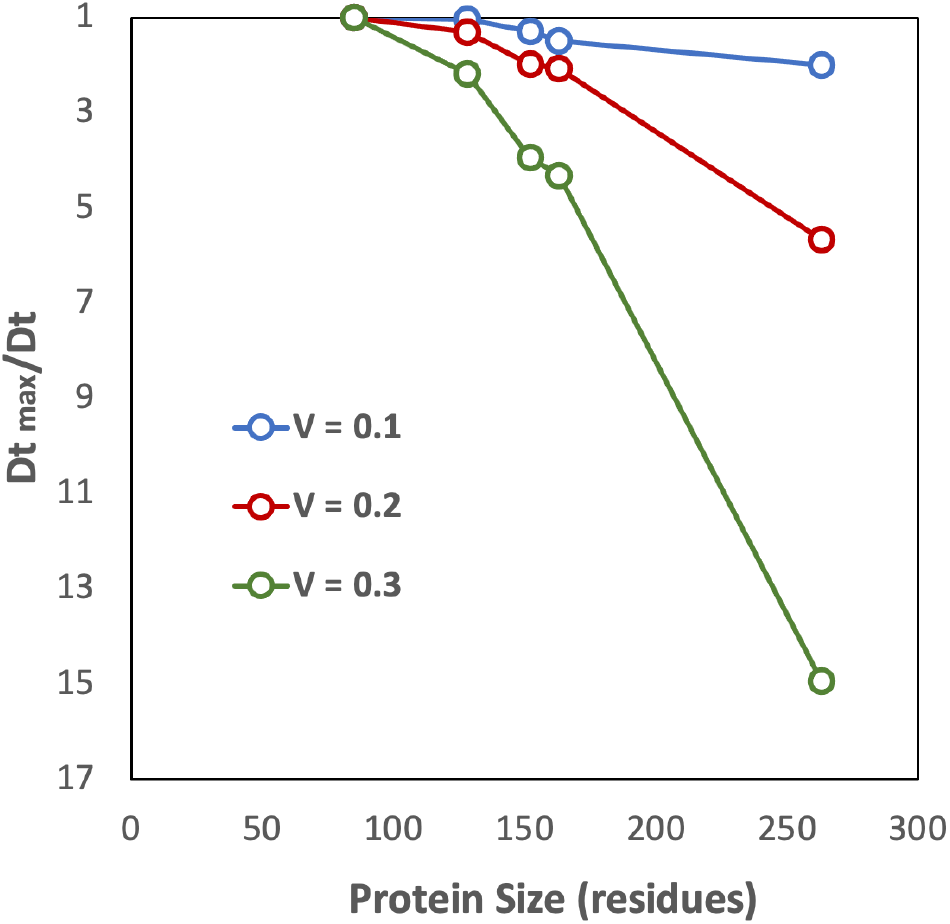
Diffusion rates vs. size of proteins. Results obtained on the “5 mix” set for the range of volume fractions. The vertical axis shows the slowdown of the diffusion rate relative to the fastest diffusion rate. The slowdown correlates with the size of the protein at all volume fractions. The rate of the slowdown increases with the volume fraction.

#### Aggregation

Experimental data on aggregation of proteins (cluster formation) at close to physiological concentrations points to the aggregation number *N_c_* (the average number of proteins in protein assemblies) for lysozyme *N_c_* ≅ 5,^47^ and monoclonal antibodies *N_c_* = 4 - 6.^48^ Our data obtained on the “5 mix” set (Fig. 8A), at the physiological volume fraction 0.3, yielded the aggregation number (cluster size) *N_c_* = 3.9, in excellent agreement with these estimates. The results show that the aggregation number does not change much across the whole range of the volume fractions (Fig. 8A). This explains similarity of the energy values *E* per molecule at different volume fractions (Figs. 3 and S3), since according to our move set, this energy is determined by the number of the protein’s interfaces with other proteins.

**Figure 8.**
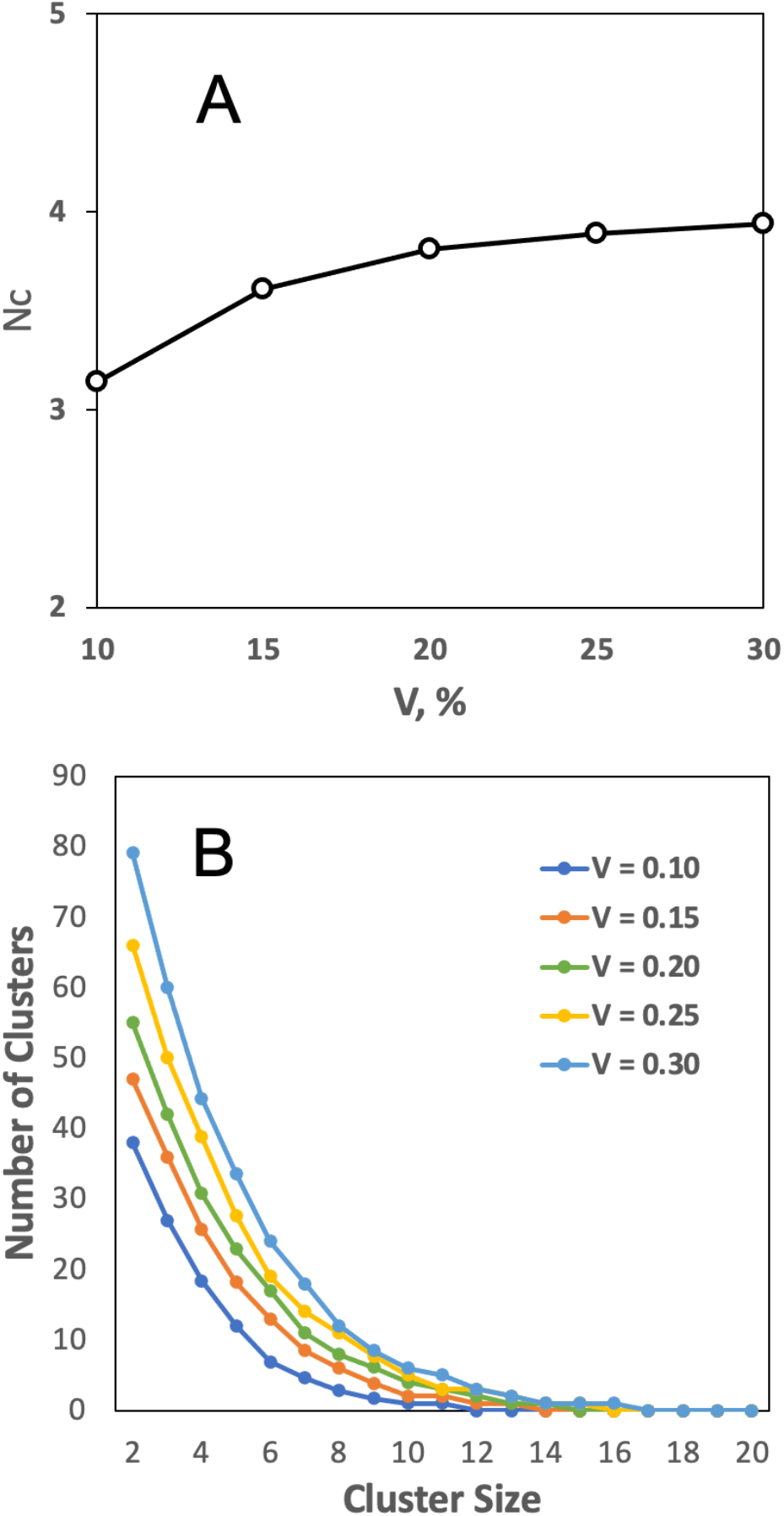
Cluster formation. (A) The aggregation number *N_c_* (the average size of protein clusters) across volume fractions *V*. (B) Distribution of cluster sizes at different volume fractions. The total number of proteins in the simulation box grows with increase of the volume fraction (Table 2). Thus, the absolute numbers of clusters at higher volume fractions are larger than those at the lower volume fractions.

The distribution of the cluster sizes (Fig. 8B) is in qualitative agreement with the results of the MD simulation in the *N_c_* = 1 - 10 range.^15^ On average, at each step of the simulation, a small percentage of proteins in our system (4% for *V* = 0.3 and 7% for *V* = 0.1) are monomers (proteins whose partners have moved away, and who have not acquired another partner yet, according to our move set).

#### Residence time

The existing estimates of the proteins’ residence time (the lifetime of a protein pair) vary dramatically among the studies. An experimental study of lysozyme protein solution determined that the protein clusters (complexes) have a lifetime longer than the time required to diffuse over a distance of a monomer diameter.^49^ Such a distance would correspond to approximately 50 steps in our simulation protocol (1 μs). The MD simulation, however, predicted far shorter lifetimes, with most times < 20 ns^13^ (one step in our simulation protocol). In our simulation, at volume fractions comparable to the ones in the above studies, ∼95% of the proteins stay in their current complex longer than one step (20 ns), but 78% move within 50 steps (i.e., move ∼1 diameter of a monomer). Thus, our results are in-between the above experimental and MD estimates.

### Trajectory length

Running the “5 mix” protein set in a 500 x 500 x 500 Å^3^ box (the smallest cell is ∼1,000 Å in linear dimension) for 10,000 steps (200 μs) at volume fraction 0.3 (Fig. 9) takes ∼5 hours on 3.1 GHz Intel Core i7 processor. The same calculation at volume fraction 0.1 takes ∼30 min. That puts a <u>0.3 - 3 second</u> simulation of such system in about one year CPU timeframe. Given the all-atom resolution of our approach, this is an extraordinary long simulation trajectory, that provides an opportunity to explicitly recreate *in silico* the physiological mechanisms that now are beyond the reach of atomic-resolution simulations.

**Figure 9.**
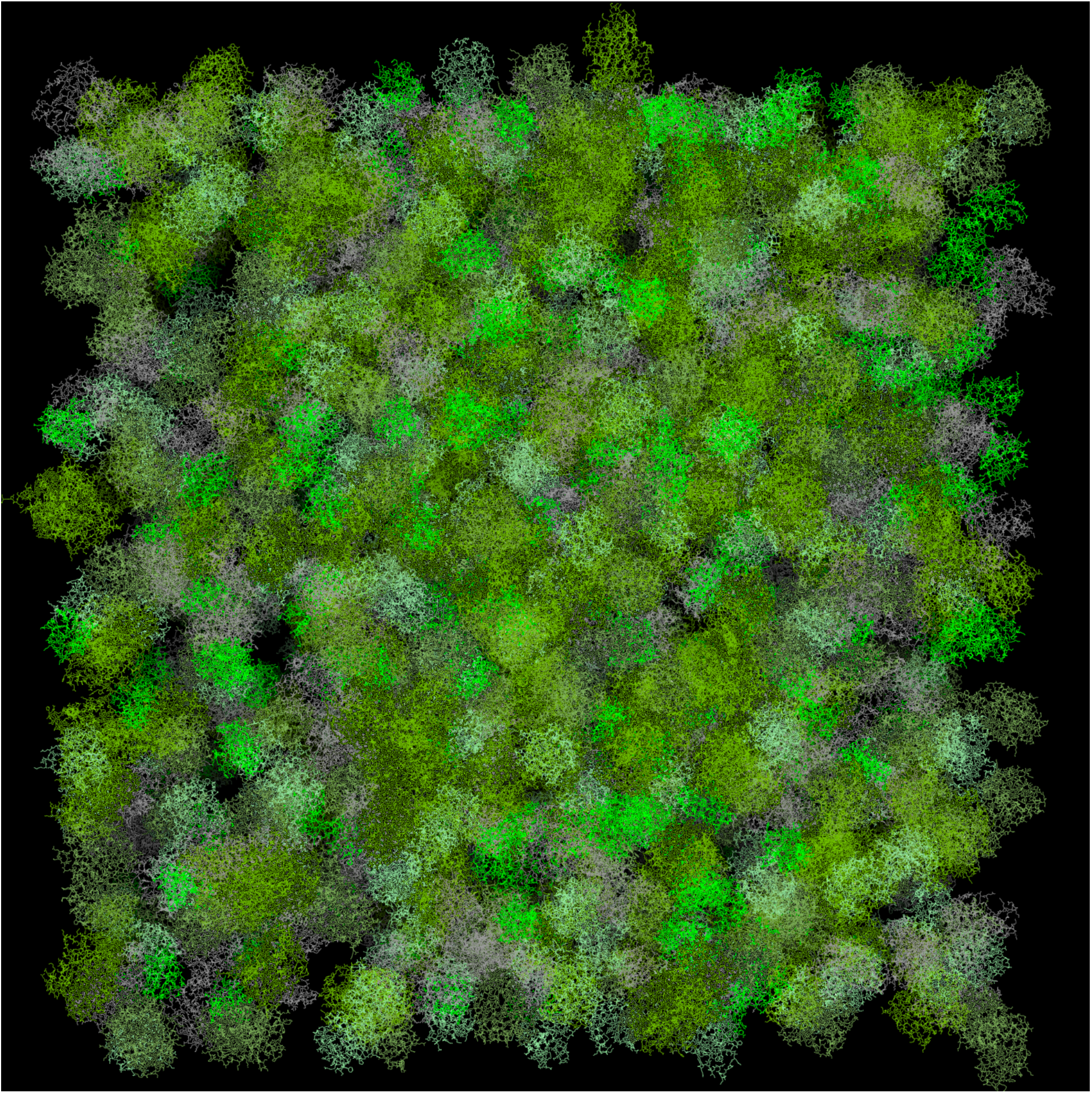
The simulation box. Protein volume fraction is the physiological 0.3. The image was obtained using PyMOL.^50^

## CONCLUSIONS AND FUTURE DIRECTIONS

Spectacular achievements of the deep learning approaches to protein structure prediction open the opportunity for protein docking to re-focus from the unique lowest energy states to the enormous multitude of the transient protein interactions that dominate the crowded cellular environment. Taking account of the transient interactions, makes it possible to propagate in time the results of static protein docking, thus taking advantage of the powerful and efficient methodologies accumulated in the protein docking field. It opens exciting opportunities in structural modeling of the protein interactions, allowing modeling of larger systems, at longer timescales, based on the atomic resolution which is integral to docking approaches.

Rapid progress in experiment and modeling is leading to the merger of molecular and cellular biology fields. New computational methodologies increasingly address modeling of the whole cell at the molecular level. The whole cell modeling can provide better understanding of cellular mechanisms and increase our ability to modulate them. The overarching goal, however, is the intellectual challenge of modeling life *in silico*.

Proteins and their interactions are the key component of cellular processes. Techniques for modeling protein interactions include protein docking and molecular simulation. The latter approaches account for the dynamics of the interactions. However, they are relatively slow, if carried out at the all-atom resolution, or significantly coarse-grained (e.g., one particle representing a protein). Protein docking algorithms (such as systematic docking by FFT) are far more efficient in sampling the spatial coordinates. However, they do not account for the kinetics of the association (i.e., do not involve the time coordinate). The approach put forward in this study bridges the two modeling techniques. The global intermolecular energy landscape of a large system of proteins was mapped by the pairwise FFT docking and sampled in space and time using MC simulations. The approach is capable of reaching unprecedented simulation timescales at all-atom resolution.

The simulation protocol was parametrized on existing MD data and validated on observations from experiments and MD simulations. The simulation performed consistently across very different systems of proteins, at a broad range of concentrations. It recapitulated data on the previously observed protein diffusion rates and aggregation. The speed of calculation allows reaching second-long trajectories of protein systems that approach the size of the cells, at atomic resolution.

This proof-of concept study is obviously just the very beginning of an expansive task of incorporating other types of macromolecules, employing more sophisticated force fields, more accurately accounting for energy barriers, introducing structural flexibility, adding membrane environment and other cellular components, multiscale modeling, and improving computational efficiency. Nonetheless, our study shows that approaches grounded in protein docking can produce unprecedented dynamic simulations of protein systems at the cellular scale.

## ACKNOWLEDGMENTS

This study was supported by NIH grant R01GM074255 and NSF grant DBI1917263. The authors wish to acknowledge contribution of Taras Dauzhenka at the early stage of this project.

## Notes

### Competing Interest Statement

The authors have declared no competing interest.

## REFERENCES

1. Jumper J, Evans R, Pritzel A, et al. Highly accurate protein structure prediction with AlphaFold. Nature. 2021;596:583–589.

2. Evans R, O’Neill M, Pritzel A, et al. Protein complex prediction with AlphaFold-Multimer. bioRxiv. 2021;doi:org/10.1101/2021.10.04.463034.

3. Vakser IA. Protein-protein docking: From interaction to interactome. Biophys J. 2014;107:1785–1793.

4. Katchalski-Katzir E, Shariv I, Eisenstein M, Friesem AA, Aflalo C, Vakser IA. Molecular surface recognition: Determination of geometric fit between proteins and their ligands by correlation techniques. Proc Natl Acad Sci USA. 1992;89:2195–2199.

5. Lensink MF, Nadzirin N, Velankar S, Wodak SJ. Modeling protein-protein, protein-peptide, and protein-oligosaccharide complexes: CAPRI 7th edition. Proteins. 2020;88:916–938.

6. Pan AC, Jacobson D, Yatsenko K, Sritharan D, Weinreich TM, Shaw DE. Atomic-level characterization of protein-protein association. Proc Natl Acad Sci USA. 2019;116:4244–4249.

7. Yu W, Jo S, Lakkaraju SK, Weber DJ, MacKerell AD. Exploring protein-protein interactions using the site-identification by ligand competitive saturation methodology. Proteins. 2019;87:289–301.

8. Li X, Moal IH, Bates PA. Detection and refinement of encounter complexes for protein– protein docking: Taking account of macromolecular crowding. Proteins. 2010;78:3189–3196.

9. Kozakov D, Li K, Hall DR, et al. Encounter complexes and dimensionality reduction in protein–protein association. eLife. 2014;3:e01370.

10. Vakser IA, Deeds EJ. Computational approaches to macromolecular interactions in the cell. Curr Opin Struct Biol. 2019;55:59–65.

11. Feig M, Sugita Y. Whole-cell models and simulations in molecular detail. Annu Rev Cell Dev Biol. 2019;35:191–211.

12. Heo L, Sugita Y, Feig M. Protein assembly and crowding simulations. Curr Opin Struct Biol. 2022;73:102340.

13. von Bulow S, Siggel M, Linke M, Hummer G. Dynamic cluster formation determines viscosity and diffusion in dense protein solutions. Proc Natl Acad Sci USA. 2019;116:9843–9852.

14. Bicout DJ, Field MJ. Stochastic dynamics simulations of macromolecular diffusion in a model of the cytoplasm of Escherichia coli. J Phys Chem. 1996;100:2489–2497.

15. Nawrocki G, Im W, Sugita Y, Feig M. Clustering and dynamics of crowded proteins near membranes and their influence on membrane bending. Proc Natl Acad Sci USA. 2019;116:24562–24567.

16. Carrera J, Covert MW. Why build whole-cell models? Trends Cell Biol. 2015;25:719–722.

17. Im W, Liang J, Olson A, Zhou HX, Vajda S, Vakser IA. Challenges in structural approaches to cell modeling. J Mol Biol. 2016;428:2943–2964.

18. Thornburg ZR, Bianchi DM, Brier TA, et al. Fundamental behaviors emerge from simulations of a living minimal cell. Cell. 2022;185:345–360.

19. Ridgway D, Broderick G, Lopez-Campistrous A, et al. Coarse-grained molecular simulation of diffusion and reaction kinetics in a crowded virtual cytoplasm. Biophys J. 2008;94:3748–3759.

20. McGuffee SR, Elcock AH. Diffusion, crowding & protein stability in a dynamic molecular model of the bacterial cytoplasm. PLoS Comp Biol. 2010;6:e1000694.

21. Wang Q, Cheung MS. A physics-based approach of coarse-graining the cytoplasm of Escherichia coli (CGCYTO). Biophys J. 2012;102:2353–2361.

22. Yu I, Mori T, Ando T, et al. Biomolecular interactions modulate macromolecular structure and dynamics in atomistic model of a bacterial cytoplasm. eLife. 2016;5:e19274.

23. Rickard MM, Zhang Y, Gruebele M, Pogorelov TV. In-cell protein-protein contacts: Transient interactions in the crowd. J Phys Chem Lett. 2019;10:5667–5673.

24. Gopan G, Gruebele M, Rickard M. In-cell protein landscapes: Making the match between theory, simulation and experiment. Curr Opin Struct Biol. 2021;66:163–169.

25. Rickard MM, Zhang Y, Pogorelov TV, Gruebele M. Crowding, sticking, and partial folding of GTT WW domain in a small cytoplasm model. J Phys Chem B. 2020;124:4732–4740.

26. Chow E, Skolnick J. Effects of confinement on models of intracellular macromolecular dynamics. Proc Natl Acad Sci USA. 2015;112:14846–14851.

27. Dill KA, Ghosh K, Schmit JD. Physical limits of cells and proteomes. Proc Natl Acad Sci USA. 2011;108:17876–17882.

28. Johnson GT, Autin L, Al-Alusi M, Goodsell DS, Sanner MF, Olson AJ. cellPACK: A virtual mesoscope to model and visualize structural systems biology. Nature Methods. 2015;12:85–91.

29. Maritan M, Autin L, Karr J, Covert MW, Olson AJ, Goodsell DS. Building structural models of a whole Mycoplasma cell. J Mol Biol. 2022;434:167351.

30. Qin S, Zhou HX. FFT-based method for modeling protein folding and binding under crowding: Benchmarking on ellipsoidal and all-atom crowders. J Chem Theory Comput. 2013;9:4633−4643.

31. Qin S, Zhou HX. Further development of the FFT-based method for atomistic modeling of protein folding and binding under crowding: Optimization of accuracy and speed. J Chem Theory Comput. 2014;10:2824−2835.

32. Nguyen TH, Zhou HX, Minh DDL. Using the Fast Fourier Transform in binding free energy calculations. J Comput Chem. 2018;39:621–636.

33. Hofling F, Franosch T. Anomalous transport in the crowded world of biological cells. Reports Prog Phys. 2013;76:46602.

34. Dix JA, Verkman AS. Crowding effects on diffusion in solutions and cells. Ann Rev Bioph. 2008;37:247–263.

35. Cruz C, Chinesta F, Regnier G. Review on the Brownian dynamics simulation of bead-rod-spring models encountered in computational rheology. Arch Comput Methods Eng. 2012;19:227–259.

36. Sanz E, Marenduzzo D. Dynamic Monte Carlo versus Brownian dynamics: A comparison for self-diffusion and crystallization in colloidal fluids. J Chem Phys. 2010;132:194102.

37. Galaktionov SG, Tseitin VM, Vakser IA, Prokhorchik YV. Amphiphilic properties of angiotensin and its fragments. Biophysics. 1988;33:595–598.

38. Kirys T, Ruvinsky A, Tuzikov AV, Vakser IA. Rotamer libraries and probabilities of transition between rotamers for the side chains in protein-protein binding. Proteins. 2012;80:2089– 2098.

39. Dauzhenka T, Kundrotas PJ, Vakser IA. Computational feasibility of an exhaustive search of side-chain conformations in protein-protein docking. J Comput Chem. 2018;39:2012–2021.

40. Shukla D, Hernandez, C.X., Weber JK, Pande VS. Markov State Models provide insights into dynamic modulation of protein function. Acc Chem Res. 2015;48:414–422.

41. He Z, Paul F, Roux B. A critical perspective on Markov state model treatments of protein-protein association using coarse-grained simulations. J Chem Phys. 2021;154:084101.

42. Voss NR, Gerstein M. 3V: Cavity, channel and cleft volume calculator and extractor. Nucl Acids Res. 2010;38:W555–W562.

43. Vakser IA. Long-distance potentials: An approach to the multiple-minima problem in ligand-receptor interaction. Protein Eng. 1996;9:37–41.

44. Ruvinsky AM, Vakser IA. Chasing funnels on protein-protein energy landscapes at different resolutions. Biophys J. 2008;95:2150–2159.

45. Hastings WK. Monte Carlo sampling methods using Markov chains and their applications. Biometrika. 1970;57:97–109.

46. Nenninger A, Mastroianni G, Mullineaux CW. Size dependence of protein diffusion in the cytoplasm of Escherichia coli. J Bacteriol. 2010;192:4535–4540.

47. Stradner A, Sedgwick H, Cardinaux F, Poon WCK, Egelhaaf SU, Schurtenberger P. Equilibrium cluster formation in concentrated protein solutions and colloids. Nature. 2004;432:492–495.

48. Scherer TM, Liu j, Shire SJ, Minton AP. Intermolecular interactions of IgG1 monoclonal antibodies at high concentrations characterized by light scattering. J Phys Chem B. 2010;114:12948–12957.

49. Liu Y, Porcar L, Chen J, et al. Lysozyme protein solution with an intermediate range order structure. J Phys Chem B. 2011;115:7238–7247.

50. The PyMOL Molecular Graphics System, Version 2.5, Schrödinger, LLC.

