## Supplementary Information for "Docking-based long timescale simulation of cell-size protein systems at atomic resolution"

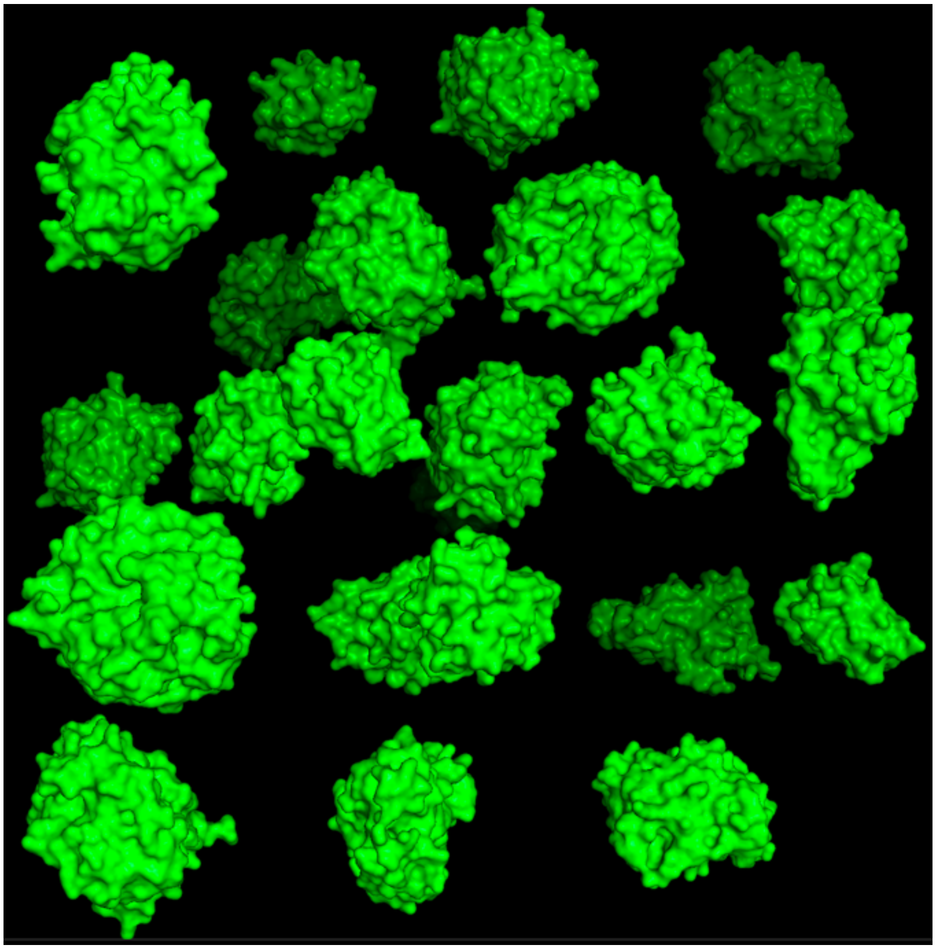

**Figure S1**. *A fragment of the initial state of the system before the start of simulation*. The volume fraction shown is 0.10. Proteins were placed on a cubical grid in random order, and randomly rotated and translated within half of the grid step. No collision check was applied since the collisions are eliminated at the start of the simulation.

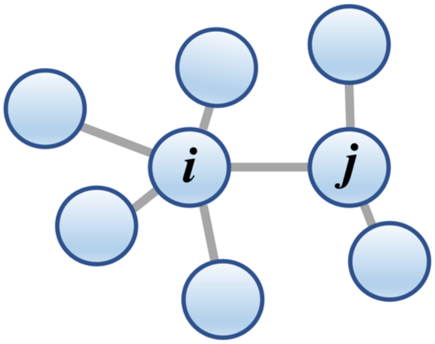


**Figure S2.** *A possible move set from states i and j.*

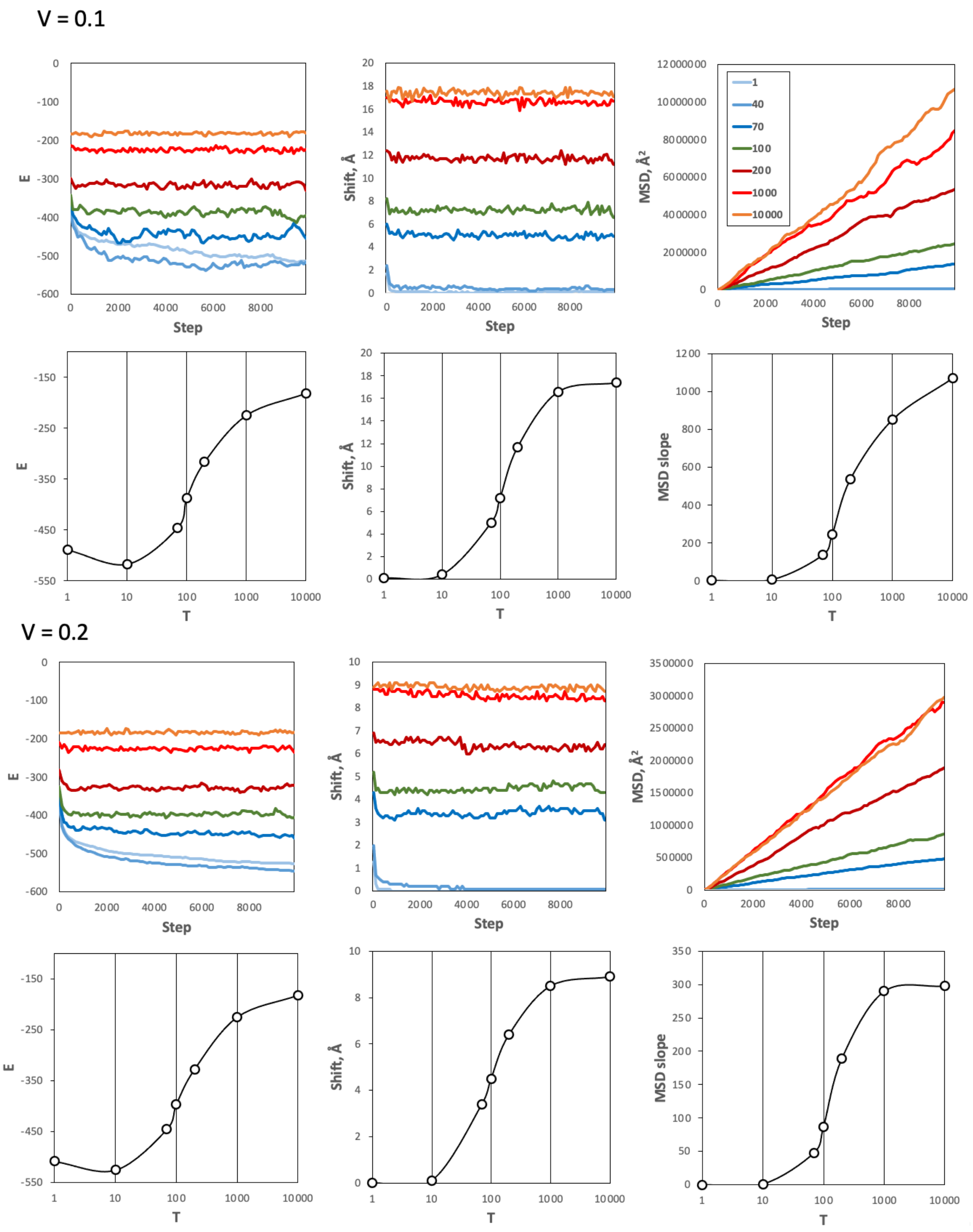


**Figure S3.** *Simulations of the "5 mix" set at lower than physiological volume fractions and a range of temperatures*. For each volume fraction *V*, the top panels show the energy *E*, shift, and MSD vs. simulation steps. MSD was calculated as the average for 1mat proteins. The temperatures *T* = 1 - 10,000 are shown by different colors. The data on the plots was smoothed by a 100-steps averaging sliding window. At low temperatures, the system is frozen (little or no movement of the proteins). At high temperatures, the system is overheated (moves accepted regardless of the energy). The melting curves (the bottom panels in log scale) have a clear inflection point at *T* = 100 indicating the optimal temperature at which the system melts (breaks from the freeze) but is not overheated yet.

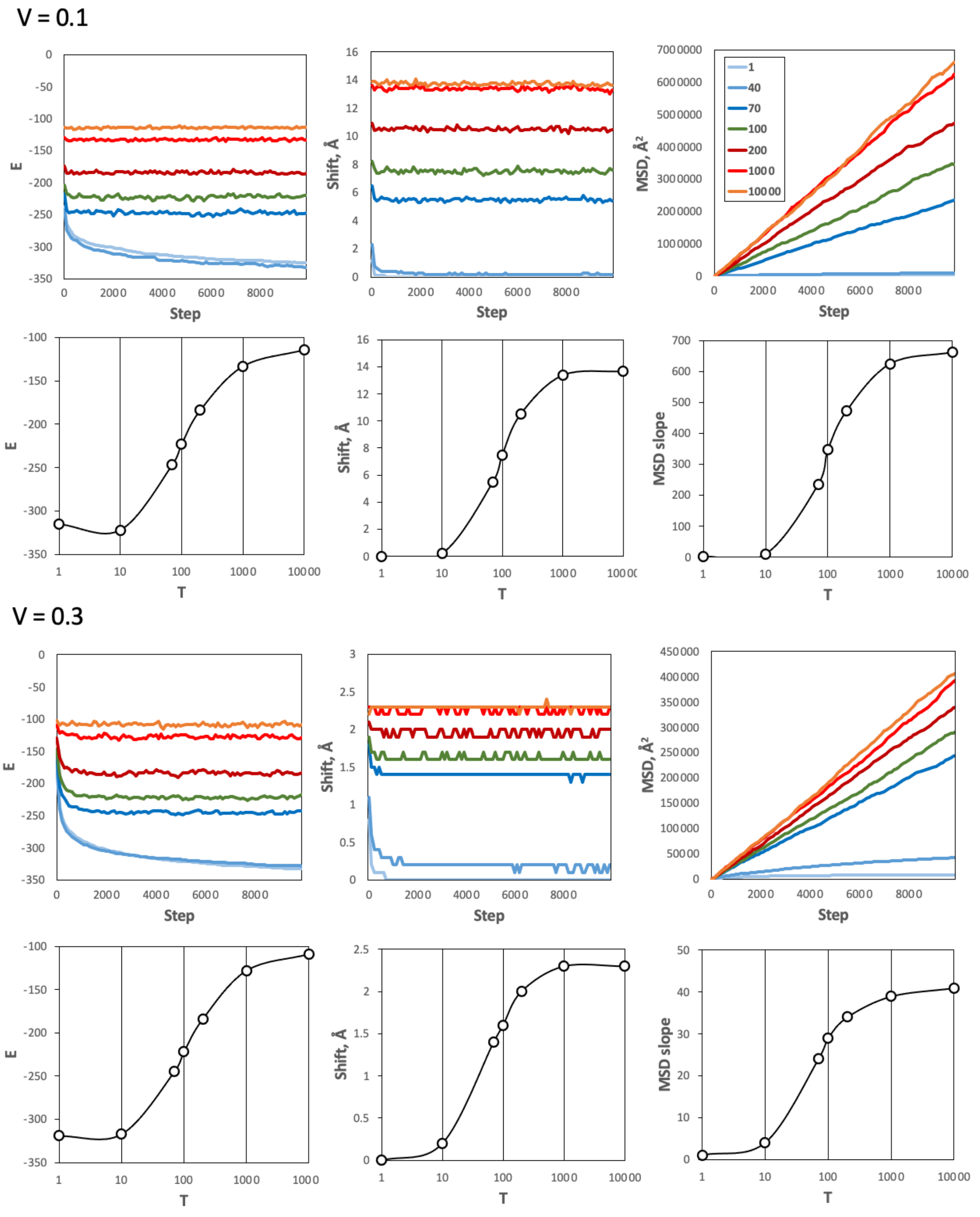


**Figure S4**. *Simulations of the "3 mix" set at low and physiological volume fractions and a range of temperatures*. For each volume fraction *V*, the top panels show the energy *E*, shift, and MSD vs. simulation steps. MSD was calculated as the average for the ubiquitin (1ubq) proteins. The details of the observable parameters are the same as in Figure S3.

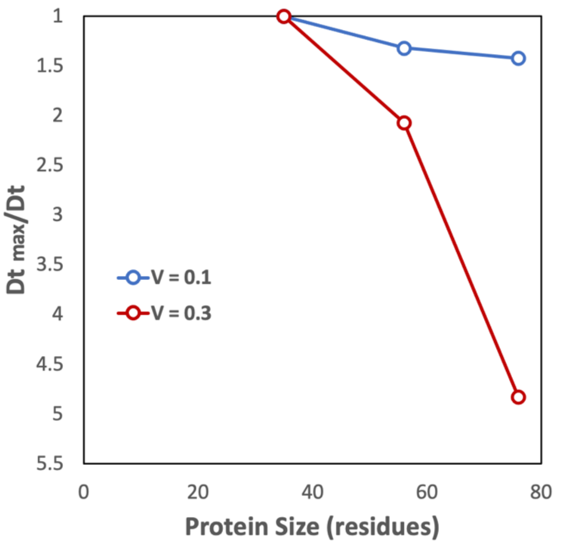


**Figure S5**. *Diffusion rates vs. size of proteins*. Results obtained on the "3 mix" set for volume fractions V = 0.1 and 0.3. The vertical axis shows the slowdown of the diffusion rate relative to the fastest diffusion rate. The slowdown correlates with the size of the protein at both volume fractions. The rate of the slowdown at the larger volume fraction is greater than that at the smaller one.
